# Microhaplotypes Improve Kinship Estimation in Heterozygous, Mixed-Ploidy Populations of *Actinidia*

**DOI:** 10.64898/2026.08.04.742852

**Authors:** Timothy Millar, Emily Koot, Astra Heywood, Adrian Grande, Susan Thomson, John McCallum, Phillip Wilcox, Michael Black

## Abstract

Over the past decade there has been increasing interest in the use of microhaplotype markers in autopolyploid taxa. This has been driven by theoretical and observed improvements in signals of allelic dosage, linkage, and heritability. Yet, to date there has been little investigation into the suitability of microhaplotype markers for estimating kinship. Here, we develop the theory of kinship estimation from microhaplotypes, introduce the MCHap microhaplotype caller for autopolyploid populations, and apply these methods to a highly diverse germplasm population of mixed-ploidy *Actinidia* (kiwifruit and relatives). We find that microhaplotype-based kinship estimates are generally superior to equivalent single nucleotide variant based estimates. This is because microhaplotypes minimize the coalescent signal among alleles which may bias estimates within the context of a recent reference population. Hence, kinship estimates from microhaplotypes more accurately capture the recent demographic history of a population. These findings are supported by both coalescent simulations and the analysis of real data. Our findings are relevant to organisms of any ploidy, but most actionable in highly heterozygous taxa such as *Actinidia*.

## Introduction

Two homologous alleles are said to be identical by descent (IBD) if either is derived from the other (or both are derived from a third) without mutation (Cotterman, 1940; Malécot, 1969). IBD is always relative to some reference point which may be defined in terms of an ancestral population or a point in coalescent time (Thompson, 2013). Kinship is the probability that randomly sampling an allele from each of two individuals — at a single locus — would yield a pair of IBD alleles (Malécot, 1969). Kinship is fundamentally distinct from phylogenetic measures of relatedness in that it considers only the proportion of shared identical alleles and does not consider the (phylogenetic) distance between non-identical alleles. This amounts to a treatment of non-IBD alleles as being equidistant. The binary nature of IBD leads to a finite set of possible relationships between any two genotypes at a given locus, as summarized by Jacquard’s identity coefficients (Jacquard, 1972) shown in Figure 1.

**Fig. 1.**
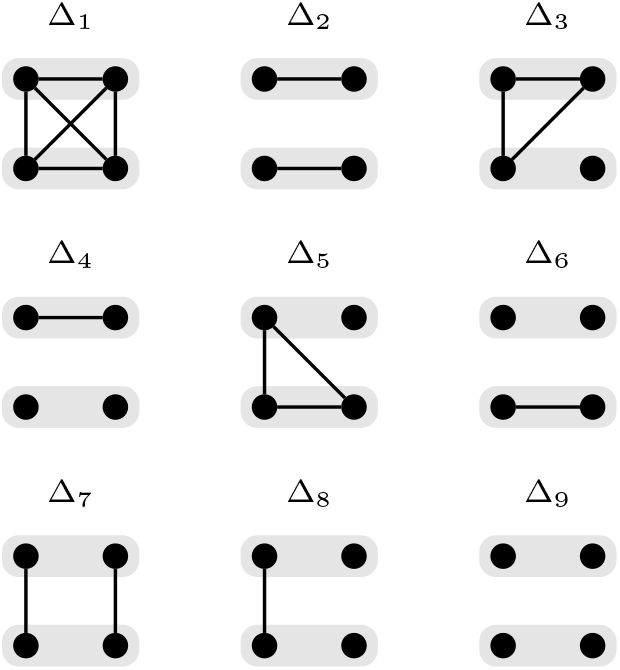
The nine condensed identity states for a pair of diploid genotypes adapted from Jacquard (1972). Within each identity, a pair genotypes are represented by filled grey areas, and the alleles of those genotypes are indicated by black dots. Lines linking alleles indicate that they are IBD.

Kinship estimators are typically defined or implemented in terms of biallelic markers such as single nucleotide variants (SNVs). This is probably due to a combination of factors including the widespread usage of SNV arrays, rarity of multi-allelic markers in commonly studied species (e.g., humans), and the inherent simplicity of working with binary data. However, the lack of allelic diversity in biallelic variants has been shown to reduce the precision of some metrics relative to that of multi-allelic markers (Hardy, 2015; Bourke et al., 2018; Thérèse Navarro et al., 2022). Furthermore, accurate dosage calling of biallelic SNVs is difficult in autopolyploids and has been identified as a source of genotyping error which can further decrease accuracy of downstream analyses (Dufresne et al., 2014; Clevenger et al., 2015; Cooke et al., 2021).

Within the past decade there have been significant advances in the development and application of targeted sequencing technologies in highly heterozygous autopolyploid crops including kiwifruit (Tahir et al., 2020), alfalfa (Zhao et al., 2023), blueberry (Clare et al., 2024; Zhao et al., 2024b), potato, (Endelman et al., 2024) and sweet potato (Zhao et al., 2024a). These are commonly ‘mid-density’ panels targeting thousands to tens of thousands of genomic loci, resulting in relatively sparse marker data compared with high-density SNV arrays. Owing to the high heterozygosity of these crops, a single sequencing target may cover multiple simple variants resulting in a high density of ‘local’ markers that can be assumed to be in complete linkage disequilibrium (LD). This results in a highly variable LD landscape among markers when calling simple variants. LD among markers is known to reduce the accuracy of kinship estimation, resulting in the development of several estimators that incorporate linkage information to improve accuracy (Wang et al., 2017).

The last decade has also seen substantial research into microhaplotype markers in diploid and polyploid species (Motazedi et al., 2017; Bourke et al., 2018; Thérèse Navarro et al., 2022; Voorrips and Tumino, 2022; Vexler et al., 2025). Microhaplotypes are defined as short haplotype sequences that can fit within the length of a single Illumina read (typically less than 200 bp) (Kidd et al., 2013). Microhaplotype markers are most informative in highly heterozygous taxa where multiple simple variants can be phased together, resulting in a highly multi-allelic marker. Multi-allelism increases the discriminatory power of markers, thereby improving the signals of linkage and trait heritability (Bourke et al., 2018; Thérèse Navarro et al., 2022; Voorrips and Tumino, 2022). Multi-allelism may also improve the accuracy of allelic dosage estimation by reducing the number of plausible dosage combinations (e.g., there can be no dosage uncertainty in the genotype ABCD). Vexler et al. (2025) proposed that relatedness estimates from microhaplotypes would emphasize recent demographic history which is consistent with the established literature on gene and nucleotide diversity (Nei, 1973; Nei and Li, 1979; Tajima, 1983). The obvious synergy between microhaplotype markers and targeted sequencing platforms represents an opportunity for the development of new statistical and software tools to aid in the analysis of autopolyploid crops.

*Actinidia* is a taxonomically complex genus of dioecious vines and scramblers found across east Asia with variable ploidy. Efforts to correctly treat the group are hampered by highly variable morphology, multiple ploidy races within taxa, and the prevalence of reticulate evolution and recurrent hybridization of taxa (Liu et al., 2017). Polyploid races of *Actinidia* are generally assumed to be autopolyploids, especially the commercially significant variates of *Actinidia chinensis* (kiwifruit). However, recent evidence indicates that some lesser-studied *Actinidia* taxa may be of allopolyploid origin including *A. valvata* (Hu et al., 2025). High heterozygosity has been reported from *Actinidia* spp. selections using a variety of marker types (Huang et al., 2004; Zhen et al., 2004; Liu et al., 2010). High heterozygosity is likely to be the result of several contributing factors including the maintenance of heterozygosity by dioecy (Ferguson and Huang, 2007), polyploidy (Udall and Wendel, 2006; Dar et al., 2017), recurrent hybridization between taxa and ploidy races (Liu et al., 2010, 2017), and large effective population size (Liu et al., 2017).

We investigate the application of microhaplotype markers for estimating kinship via theoretical development and simulation. We then introduce the MCHap software package for assembling microhaplotypes in mixed-ploidy populations. Finally, we apply our methods to a highly heterozygous *Actinidia* germplasm collection with variable ploidy levels using a Capture-GBS targeted sequencing panel. Our central hypothesis is that microhaplotypes improve the accuracy of kinship estimates with respect to a recent reference population.

## Materials and Methods

### Kinship estimation

Weir and Goudet (2017) introduced a kinship estimator based on allelic matching among pairs of samples. They first estimate the matching coefficient, which is defined as the probability that randomly drawn alleles from each of two individuals at a random locus are identical by state (IBS). The matching coefficient is then rescaled to an estimate of kinship using the expected matching coefficient of a reference population. In lieu of a known reference point, they define the coefficient *β* as the kinship scaled such that the mean estimate among samples is zero. This is estimated from the matching coefficient as:

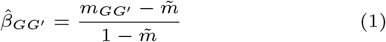

where 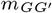is the matching coefficient between a pair of genotypes *G* and *G*′, and 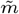 is the mean matching coefficient among all pairs of genotypes in the sample population (excluding self-matching). Within this manuscript, we focus exclusively on the Weir-Goudet approach to estimating kinship. This is motivated by its clear interpretation, straightforward generalization to both autopolyploid and multi-allelic data, and prior evidence of its accuracy within closely related populations (Goudet et al., 2018; Bilton, 2020). Furthermore, their approach does not rely on ancestral allele frequencies, which can deviate substantially from sample population allele frequencies in the presence of selection or inbreeding (VanRaden, 2008).

The matching coefficient between a pair of genotypes would typically be calculated relative to the total number of SNVs found in a sample population:

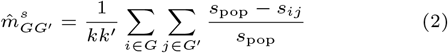

where *s*_pop_ is the total number of SNVs found in the sample population, *s*_*ij*_ is the number of SNVs differentiating *i*^th^ haplotype of *G* from *j*^th^ haplotype *G*′, and *k* and *k*′ are the ploidy of *G* and *G*′ respectively (Bilton, 2020). However, the use of *s*_pop_ is somewhat arbitrary, and an artifact of SNV discovery that depends upon the wider sample population. The *s*_pop_ term does not influence the estimate of *β*, as it factors out when substituting equation 2 into equation 1 (see Supplementary Text S1). We instead replace *s*_pop_ with the total number of base positions from which SNVs have been discovered (*n*), such that:

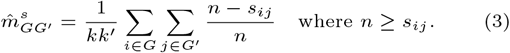

This results in a consistent matching coefficient between any pair of individuals, irrespective of the wider sample population. The value of *n* may refer to the genome length in the case of whole-genome resequencing, or the total length of sequencing targets in the case of targeted sequencing panels. Alternatively, *n* may indicate the length of a single target locus when reasoning about single locus estimates.

### Single locus expectations

Let *δ*_*ij*_ denote the evolutionary distance between the pair of alleles *i* and *j* at a single locus (i.e., twice the number of generations occurring since their most recent common ancestor). The expected number of mutations occurring between *i* and *j* is:

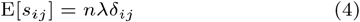

where *λ* is the mutation rate and *n* is the length of the locus in base positions. For simplicity, we ignore both recombination and the probability of multiple mutations occurring at a single base position. The expected matching coefficient between a pair of sequences when calculated from IBS of SNVs is the mean of probabilities that no mutations have occurred at each base position:

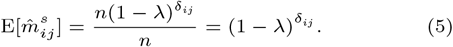

which follows an exponential distribution when viewed as a function of evolutionary time. The expected matching coefficient between a pair of genotypes at this locus is:

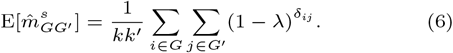

Consider a set of allelic states resulting from a coalescent tree. Observations of simple mutations such as SNVs within each allele are partial observations of the full allelic state (i.e., the full sequence). A single biallelic marker partitions the set of realized alleles into a pair of subsets that cannot be IBD. The combination of all mutations across a genomic segment partitions the realized alleles into IBS sets that each correspond to one or more IBD subsets with respect to some reference point. In practice we cannot distinguish pairs of alleles that are IBD from those that are merely IBS (Jacquard, 1972). We instead follow the approach of Weir and Goudet (2017) in which probabilities of IBD are estimated by rescaling probabilities of IBS. Given that non-IBS necessitates non-IBD (Cotterman, 1940; Malécot, 1969), we aim to improve these estimates by refining the signal of non-IBD among observed allelic states.

Any pair of mutations on a single coalescent tree will encode some amount of redundant information about IBS, and thereby IBD. If both mutations occur on the same branch of the tree, they encode identical partitions of the alleles and are fully redundant. A pair of biallelic mutations occurring on different branches encode different partitions that must jointly define three subsets of alleles. However, the alleles corresponding to two of these subsets can be differentiated by both mutations. This will result in a greater mean of estimates of non-IBS between these alleles, relative to that of allele pairs that are differentiated by only a single variant. The increased weighting given to the difference between these alleles is a violation of the assumption of equidistance among non-IBD alleles, and will translate into variation in kinship estimates among genotypes containing these alleles.

Under a bifurcating model of sequence evolution, it is improbable that all combinations of non-IBD alleles will have identical (expected) matching coefficients when calculated from SNV markers (Tajima, 1983). The example in Figure 2A illustrates the evolution of three non-IBD alleles, of which *a* and *b* are more recently diverged than *c*. In this scenario we would *expect* that there are fewer SNVs differentiating alleles *a* from *b* than there are differentiating *c* from either *a* or *b*. Hence, the expected matching coefficient of genotypes drawn from these alleles will be influenced by the coalescent history of alleles in addition to the identity state of the genotypes. This can result in different expected matching coefficients for allelic permutations of a single identity coefficient, as shown for Δ_4_ in Figure 2B. This observation implicates coalescent history as a source of bias in kinship estimation and motivates the development of methods that minimize the coalescent signal among non-IBD alleles.

**Fig. 2.**
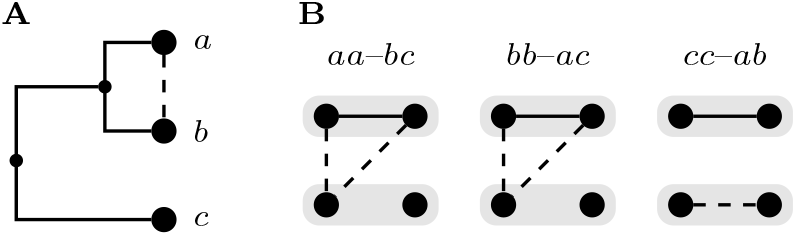
**A**: Bifurcated coalescent tree of three alleles *a, b* and *c*. **B**: Alternate allelic combinations drawn from **A** resulting in genotype pairs corresponding to Jacquard’s identity coefficient Δ_4_. Solid lines indicate IBD alleles as in Figure 1. Dashed lines indicate the allele pair *a*–*b*, which are expected to have higher sequence similarity than either *a*–*c* or *b*–*c*.

Consider a small genomic locus of length *n* ≲ 200 base positions. Sequences from such a locus are microhaplotype alleles, which may comprise multiple SNVs. Two such microhaplotype alleles are IBS if their entire sequences are identical. This leads to the haplotype matching coefficient that has an expected value of:

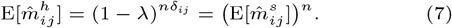

Hence, the distinction between the microhaplotype and SNV matching coefficients is analogous to the distinction between Nei’s gene and nucleotide diversities (Nei, 1973; Nei and Li, 1979). The haplotype matching coefficient ensures that non-IBS alleles are treated as equidistant within the span of a given genomic locus and, in doing so, eliminates redundant information among the SNVs occurring within that locus. It follows that the mean of estimates of the microhaplotype matching coefficient should generally exhibit less influence from coalescent history than the equivalent SNV matching coefficient.

When rescaling the expected microhaplotype and SNV matching coefficients following equation 1, we find:

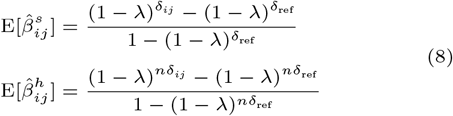

which describe a pair of negative exponential curves intersecting at 1 and 0 such that:

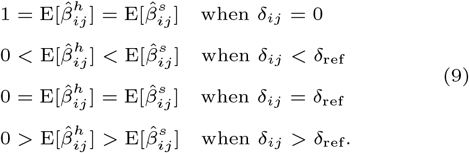

Therefore, the microhaplotype expectation of *β* between non-IBD alleles is always closer to their true kinship (zero) than the SNV-based expectation. We note that both the microhaplotype and SNV estimates will be one (i.e., congruent with IBD) when the realized alleles are in fact IBD (irrespective of *δ*_*ij*_).

The variance of the SNV and haplotype matching coefficients for a length *n* locus can be modeled as (the mean of *n*) random Bernoulli variables:

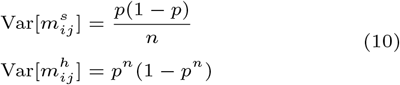

where 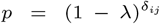 (i.e., the expectation of the SNV matching coefficient). However, these variances are not directly comparable because of the difference in expectations of the SNV and haplotype matching coefficients. We can rescale these variance terms following equation 1 by replacing 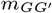with *p* and 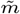 with 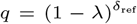 (i.e., the expected SNV matching coefficient of unrelated individuals). The variance terms are then:

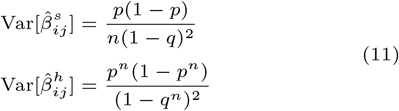

which indicates that haplotype-based estimates generally display less variance. An exception to this occurs for pairs of individuals that are closely related, relative to the population mean (i.e., when *p* is almost 1 and *q* is substantially smaller). For a pair of unrelated individuals (i.e., *p* = *q*) equation 11 simplifies to:

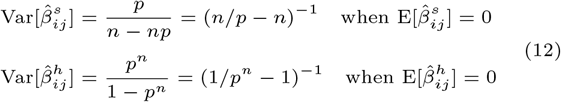

which, with constraints *n >* 1 and 0 *< p <* 1, shows that the variance of the haplotype estimate between unrelated individuals will be smaller than that of the SNV estimate.

Consider a population of diploid genotypes that share a total of three unique in descent alleles, as shown in Figure 2A. Enumerating all possible combinations of diploid genotypes reveals the seven unique expected matching coefficients shown in Table 1.

**Table 1.** Expected matching coefficients of all unique combinations of diploid genotypes in a population with three unique alleles derived as shown in Figure 2A. The possible identities (Figure 1) corresponding to each expected matching coefficient are also shown.

| ID | Genotype pairs | Identities |
| --- | --- | --- |
| $\mu_1$ | *aa-aa, bb-bb, cc-cc* | $\Delta_1$ |
| $\mu_2$ | *aa-ab, ab-ab, ab-bb* | $\Delta_3, \Delta_5, \Delta_7$ |
| $\mu_3$ | *aa-bb* | $\Delta_2$ |
| $\mu_4$ | *aa-ac, ac-ac, bb-bc, bc-bc, ac-cc, bc-cc* | $\Delta_3, \Delta_5, \Delta_7$ |
| $\mu_5$ | *ab-ac, ab-bc, ac-bc* | $\Delta_8$ |
| $\mu_6$ | *bb-ac, aa-bc* | $\Delta_4, \Delta_6$ |
| $\mu_7$ | *aa-cc, ab-cc, bb-cc* | $\Delta_2, \Delta_4, \Delta_6$ |

We can visualize the difference in expected matching coefficients as calculated from SNV or microhaplotype alleles by assuming some model parameters. In Figure 3 we observe that matching coefficients involving one or more heterozygous genotypes fall to the right of the line of possible haploid matching coefficients (coefficients *μ*_1_, *μ*_3_ and *μ*_7_ occur between pairs of homozygous genotypes and are equivalent to haploid matching coefficients). The sole exception is the genotype pair *ab*-*cc* in *μ*7, where alleles *a* and *b* are equidistant from *c*. The rightward shift of heterozygous genotypes indicates a relative increase in the expected matching coefficients of genotype pairs involving different coalescent branch lengths when calculated from microhaplotypes. Following equation 7, we can see that this must always be the case because the mean of any power of two values in the interval (0, 1) will always shift towards that of the smaller value as the exponent increases (i.e., as the locus length *n* increases). This shift can result in a rank order change of expected matching coefficients as observed for *μ*_3_ and *μ*_4_ in Figure 3. Rank order shifts in *realized* matching coefficients can occur within loci that span as few as three biallelic variants, as demonstrated in Figure 4.

**Fig. 3.**
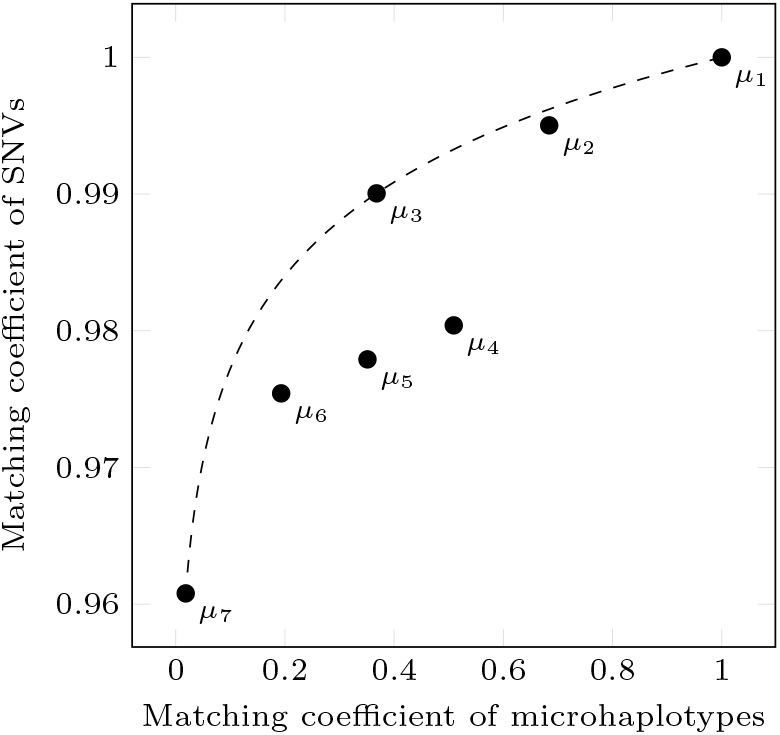
The seven unique expected diploid matching coefficients for single nucleotide variant (SNV) and microhaplotype markers as calculated when *δ*_*ab*_ = 10^4^, *δ*_(*ab*)*c*_ = 4 *×* 10^4^, *λ* = 10^*−*6^ and haplotype length of *n* = 100bp. The dashed line indicates possible expected matching coefficients between pairs of haplotypes for any value of *δ*.

**Fig. 4.**
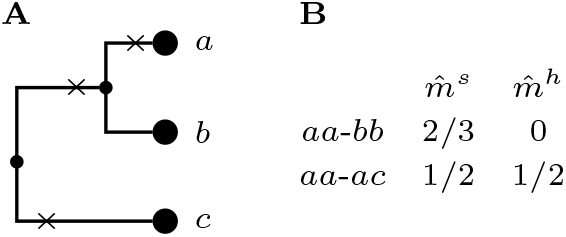
**A**: Bifurcated coalescent tree of three alleles *a, b* and *c*. The ‘*×*’ symbols indicate separate SNVs with respect to the ancestral allele. **B**: The resulting realized matching coefficients for two pairs of diploid genotypes as estimated from single nucleotide variant (SNV) 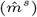 and haplotype markers 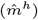.

Changes in the rank order of relationships may also occur within haploid genotypes when allowing for recombination. Consider again the three alleles *a, b* and *c* derived from a single common ancestor, as shown in Figure 2A. Now assume that these alleles correspond to a genomic region extending over a pair of equally sized microhaplotype loci. The matching coefficient between a pair of haploid individuals across both loci is calculated similarly to the matching coefficient of a pair of diploid individuals at a single locus. Hence, it is trivial to reconstruct an equivalent of Figure 3 by enumerating all possible pairs of recombinant haplotypes across both loci.

### Simulation analysis

We performed coalescent simulations in msprime (V1.3.3, Baumdicker et al., 2021) of a 22 Mbp chromosome with a genetic length of 150 cM which is comparable to the *A. chinensis* linkage group 3 described by Tahir et al. (2020). The simulated demographic history was designed to align with parameters used to obtain Figure 3 with the aim of confirming the theoretical expectations developed in the previous sections. To this end, we simulated three subpopulations A, B and C with population splits A–B and (AB)–C at 10^4^ and 4 *×* 10^4^ generations before the present, respectively. This simulation was repeated in full using less extreme evolutionary distances by specifying population splits for A–B and (AB)–C at 10^3^ and 4 *×* 10^3^ generations, respectively. For simplicity, we respectively refer to these as ‘deep’ and ‘shallow’ simulations throughout the remainder of this manuscript.

In both simulations the population sizes were held at a constant 5000 individuals for each sub-population and ancestral population. Single variant mutations were simulated using a constant mutation rate of 10^*−*6^. Finally, we simulated a single admixed generation with parents chosen randomly with replacement from sub-populations A, B and C. Ten diploid genotypes where sampled from each sub-population (A, B and C) and a further 100 were sampled from the single admixed generation. The SNV and haplotype matching coefficients were estimated from 1000 evenly spaced genomic intervals of 100bp in length. The SNV estimate of the matching coefficient was calculated by normalizing over the sum of loci lengths as in equation 3. Matching coefficient estimates were then transformed into estimates of *β* following equation 1 and visualized as a scatter plot.

The ‘true’ kinship among simulated genotypes was calculated by extracting IBD segments from the simulated data using tskit (V0.6.4, Kelleher et al., 2016; Wong et al., 2024) and calculating probabilities of IBD from the proportion of shared IBD segments across the whole chromosome (i.e., not restricted to the 1000 genomic intervals). We note that this method of identifying IBD segments considers only inheritance from the reference population (descent) and ignores mutations with respect to the ancestral allele (state). The true kinship was then transformed to *β* following equation 1 and used to estimate the mean squared error of the SNV and haplotype estimates. This process was repeated for a range of reference population depths (10^0^, 10^1^, …, 10^4^ generations) by considering only allelic inheritance since that point.

### Microhaplotype calling

We construe genotyping of polyploid individuals using haplotypes as the task of finding the optimal genotype among the set of all possible phased genotypes. Let *G* be the set of all possible phased genotypes of ploidy *k* within a genomic locus. A given genotype *G ∈ G* is a multiset of haplotype alleles with cardinality |*G*| = *k*. The posterior probability of each possible genotype can be inferred via Bayes’ Theorem as:

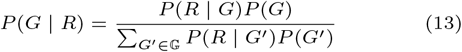

where *R* is a set of observed read sequences.

The number of possible genotypes (i.e., the cardinality of *G*) is a function of the individual’s ploidy and the number of possible haplotype alleles. Let ℍ be the set of all unique haplotypes that could occur within a genomic locus. The number of unique genotypes is found by enumerating all possible combinations of *k* haplotypes allowing for duplicated alleles, which leads to the multiset coefficient. The number of possible haplotype alleles (i.e., the cardinality of ℍ) is calculated as the product of the number of unique alleles found at each SNV position within the genomic locus. Hence, evaluation of the denominator in equation 13 is often intractable.

We employ two Markov chain Monte Carlo strategies to avoid direct enumeration over *G*. The first of these is a Metropolis-Hastings algorithm (Hastings, 1970), which is suitable for de novo assembly of SNVs into microhaplotypes using short-read data and is designed to be generalized to variable stepping criteria. Within the *t*^th^ iteration of this algorithm, the current genotype *G*^*t*^ is replaced with a new genotype *G*^*t*+1^ selected from a family of neighboring genotypes, or if all neighboring genotypes are rejected, *G*^*t*+1^ = *G*^*t*^. Given a genotype *G* and a finite family of neighboring genotypes *N* ^*t*^, the probability of a transition to each neighboring genotype (*G*′ *∈ N* ^*t*^) is calculated using the Metropolis-Hastings rule:

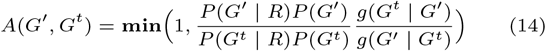

where the proposal function *g* depends on the number of neighboring genotypes:

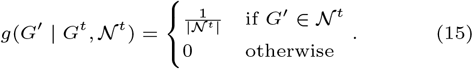

The probability of transition from *G*^*t*^ to *G*′ is then calculated as the joint probability of both proposing and accepting *G*′:

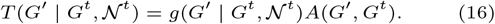

This forms the basis of a generic microhaplotype assembly algorithm that allows for any combination of neighborhood functions.

The choice of neighborhood function(s) is essential to both the validity and efficiency of the MCMC. The simplest method of identifying neighboring genotypes is to ‘mutate’ a SNV within a single haplotype of the current genotype. This is a reversible step which, when repeated, allows the Markov chain to transition between any pair of genotypes, thereby satisfying the requirement of irreducibility (Gelman et al., 2014). A common issue in autopolyploids is the possibility of multiple high probability genotypes that are dosage variations of the same set of unique haplotypes. This can result in multi-modality when relying on the ‘mutation’ function alone. To mitigate this issue, we introduce a ‘dosage’ function which identifies neighboring genotypes by altering the dosage of the current genotype as shown in Figure 5. Similarly, we introduce a ‘recombination’ function in which subsections of the haplotypes within a genotype may be exchanged with the aim of improving mixing efficiency.

**Fig. 5.**
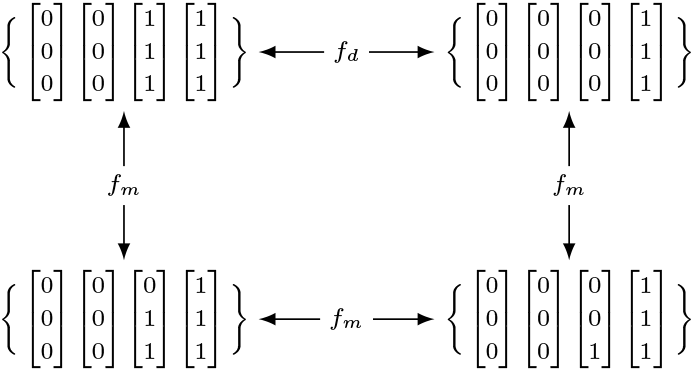
Example of a ‘dosage’ step allowing more efficient transition between tetraploid genotypes of alternate dosages. Each genotype is encoded as a multiset of haplotype vectors whose elements indicate the allelic state at each of three variable positions. Edges connecting genotypes show possible transitions between genotypes based upon mutation (*f*_*m*_) and dosage (*f*_*d*_) steps.

In addition to the Metropolis-Hastings algorithm described above, we also employ a Gibbs sampler (Geman and Geman, 1984) for calling genotypes from a set of known haplotypes. Within each step of the Gibbs sampler, a single haplotype of a genotype is updated by iterating over the set of known haplotypes to calculate the marginal distribution. This approach is preferred when the set of known haplotypes is relatively small, as it is less prone to mixing issues and allows for simpler specification of a prior distribution if desired.

Both our Metropolis-Hastings and Gibbs samplers utilize a likelihood function that is similar to those of existing tools including ANGSD (Korneliussen et al., 2014) and HapTree (Berger et al., 2014), but extended to incorporate multi-allelic SNVs. This assumes that observed read sequences are drawn from the true genotype following a multinomial distribution:

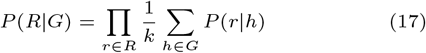

where *r* and *h* are sequences of allelic states from an observed read and proposed haplotype respectively. Sequencing errors in observed reads are accounted for with an error rate that may be constant (default of 0.0024 (Pfeiffer et al., 2018)) or informed by the reported quality scores of individual nucleotides.

Efficient implementations of these algorithms are available through our software package MCHap, which is suitable for the parallel assembly and inference of hundreds of genotypes across thousands of genomic loci. MCHap is composed of multiple sub-tools, including MCHap ‘assemble’ for *de novo* assembly of haplotypes using the Metropolis-Hastings algorithm, and MCHap ‘call’ for inferring genotypes from a set of known haplotypes via a Gibbs sampler. These tools can be applied to genotypes of any ploidy (assuming autopolyploidy), and ploidy may differ among genotypes within a single analysis. MCHap allows for the specification of a flat prior across genotypes or a Dirichlet-multinomial prior that incorporates expected allele frequencies and inbreeding coefficients following Blischak et al. (2017). MCHap makes extensive use of standard bioinformatics file formats, including BAM, BED, Fasta and VCF (Li et al., 2009; Danecek et al., 2011), and is released under the MIT software license at https://github.com/Plant-Food-Research-Open/MCHap.

### Application to kiwifruit germplasm

A sample population of 97 genotypes were identified from the germplasm collections across Kerikeri, Auckland, Te Puke, Te Papaioea (Palmerston North) and Motueka corresponding to 17 *Actinidia* taxa across five ploidy levels (see Table 2). Samples were collected over three consecutive summers (2021– 2023) with 1cm *×* 1cm sections of leaf being collected from orchards, greenhouses and tissue culture, and individually stored in 2 mL screw cap O-ring tubes half filled with silica beads. Leaves were collected in duplicates. DNA was extracted using a modified CTAB protocol (Gardiner et al., 1996). DNA was then quantified using the Quant-iT™ PicoGreen™ dsDNA Assay kit (Invitrogen, Burlington, ON, Canada) and a SpectraMax^®^ Gemini EM Microplate Reader (Molecular Devices, San Jose, California, USA) and 1 μg of DNA was provided to Rapid Genomics LLC, Gainesville, Florida, USA for Capture-sequencing. The Capture-sequencing array utilized was the same array described by Tahir et al. (2020) comprising approximately 10, 000 exonic baits targeting 120 bp loci and was originally designed from diploid *A. chinensis* var. *chinensis* data.

**Table 2.**
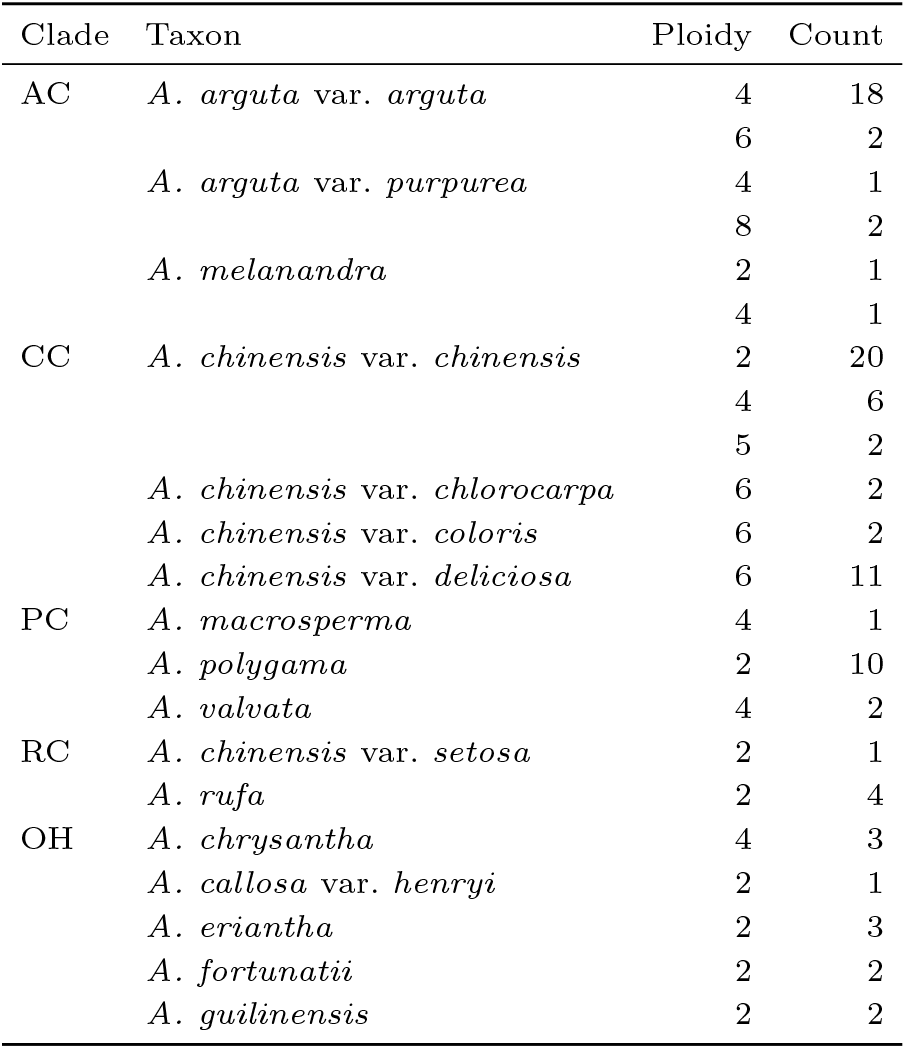
Germplasm genotypes by clade, taxon and ploidy. Clades correspond to those described by Liu et al. (2017) with AC = *Actinidia arguta* complex; CC = *A. chinensis* complex; AC = *A. polygama* clade; RC = *A. rufa* clade: OH = other hairy and/or lenticillate fruited taxa.

Read adapters and low-quality termini were removed using AdapterRemoval (V2.2.2, Schubert et al., 2016) with gzip compression and read-pair collapsing enabled to merge overlapping pairs. Reads were aligned to *Actinidia chinensis* var. ‘Donghong’ (Han et al., 2023) with BWA-MEM (V0.7.18 Li and Durbin, 2009). Paired-end reads and collapsed/singleton reads were mapped separately to preserve library structure and read-group labeling. For each read set, paired and singleton BAMs were merged and name-sorted prior to mate-information correction using Samtools (V1.20, Li et al., 2009). Each sample alignment job was performed with 2–4 CPUs and 16–32 GB RAM.

To find read intervals that were represented across the population, the cram files were processed using custom python scripts. For each sample, input CRAM files were first converted into BED format using bedtools bamtobed (V2.30.0, Quinlan and Hall, 2010), retaining chromosome, start, end, and CIGAR string information. Identical intervals were collapsed, and overlapping intervals were clustered to identify consensus regions. Within each cluster, intersection coordinates were defined by the latest start and earliest end positions, while read support was quantified by summing per-interval counts. Intervals were re-oriented to ensure consistent genomic coordinates, annotated with their length, and filtered to retain only those supported by more than 50 reads. The final per sample high-confidence intersecting intervals were aggregated, concatenated and clustered with bioframe (V0.8.0, Abdennur et al., 2024), enabling the grouping of overlapping regions into cluster-level intervals. For each cluster, the proportion of unique sample identifiers was calculated, and clusters were retained if they were represented in at least 98% of samples. Each bait site was restricted to 100 bp, with 50 bp either side of the center of the interval. The resulting coordinates were exported in BED format for haplotype assembly.

Haplotypes were called using an empirical Bayes strategy in an attempt to mitigate the extreme heterozygosity and unknown population structure of the *Actinidia* germplasm. Putative SNV loci were identified from each sample using MCHap’s ‘find-snvs’ tool based on simple thresholds including a minimum allele depth of four and individual allele frequency of 0.1. *De novo* haplotype assembly of the putative SNV loci was then performed using MCHap’s ‘assemble’ tool with a flat prior across genotypes (i.e., informed only by the likelihood function). Haplotype assembly was constrained to the 100 bp windows identified from aligned Capture-seq reads. The assembled haplotypes were filtered to exclude those with a posterior probability of *<* 0.5 of occurring within at least one individual of the sampled germplasm (using the ‘AOP’ field reported by MCHap). Genotypes were then called with a Gibbs-sampler using MCHap’s ‘call’ tool to restrict the candidate haplotypes to those identified across the sampled germplasm in the previous step. Genotype calling also utilized a flat prior. Each stage of the haplotype-calling pipeline was run in ten parallel jobs by splitting the BED file of targets into ten separate chunks. All analyses were performed with MCHap V0.11.1.

The resulting VCF file of haplotype calls was converted to a Zarr store using bio2zarr (V0.1.5, Czech et al., 2025) and then analyzed with sgkit (V0.10.0, https://github.com/sgkit-dev/sgkit). Haplotype calls were filtered in sgkit to remove any monomorphic variants and any loci with mean support quality *<* 40, where support quality is calculated as *−*10 *×* log_10_(*P* (*A*)) and *P* (*A*) is the posterior probability of the set of called alleles irrespective of their dosage (reported as ‘SQ’ by MCHap). The haplotype VCF file was also converted to SNV calls using MCHap’s ‘atomize’ tool. SNV calls were similarly converted to a Zarr store and filtered in sgkit such that equivalent samples and loci were retained. Matching and *β* coefficients were estimated from both the haplotype and SNV datasets using the functions ‘identity by state’ and ‘Weir Goudet beta’ in sgkit. We note that the implementation of ‘identity by state’ in sgkit estimates the matching coefficient following equation 2 rather than equation 3 as used for the simulation results. Principal component analysis (PCA) of matching coefficients was performed using scikit-learn (V1.7.0, Pedregosa et al., 2011). Finally, the microhaplotype- and SNV-based coefficients were visually compared via a scatter-plot.

In addition to pairwise kinship, we also estimated within taxon allelic diversity for taxa represented by at least ten samples. Within-taxon microhaplotype diversity was estimated by calculating the genome-wide mean gene diversity (Nei, 1973) of microhaplotypes using the function ‘diversity’ in sgkit. The within-taxon genome-wide mean of nucleotide diversity (Nei and Li, 1979) was calculated using the same function applied to SNVs within genomic windows corresponding to the assembled microhaplotypes.

## Results

### Coalescent simulation

Chromosome-wide matching coefficients among simulated genotypes yielded drastically different estimates when estimated from SNV and microhaplotype markers (Figure 6), consistent with theoretical expectations (Figure 3). Microhaplotype and SNV estimates were much more comparable when rescaled to estimates of *β* (alternate axes in Figure 6). Substantial evidence of rank-order variation between SNV- and microhaplotype-based estimates were present in both simulations, but most extreme in the ‘deep’ simulation.

**Fig. 6.**
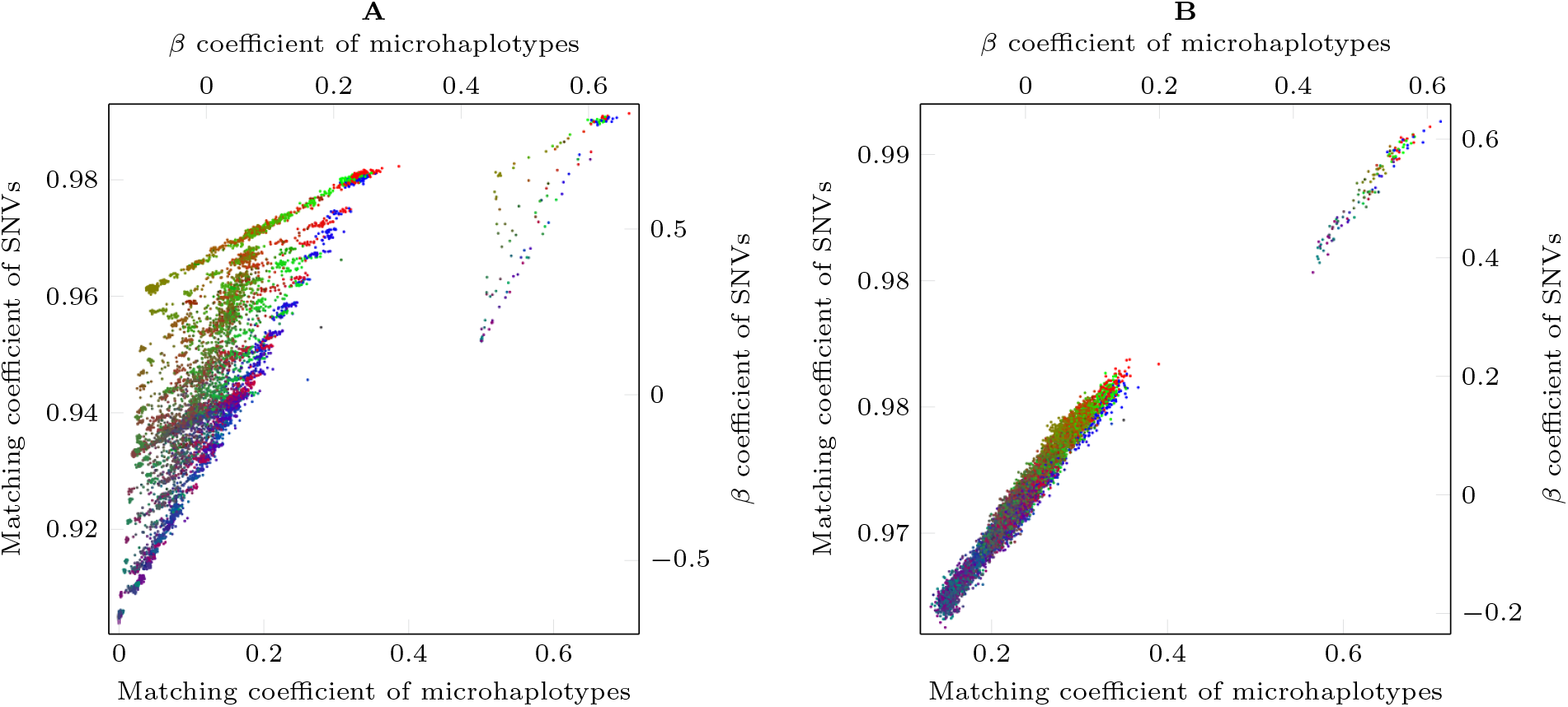
Estimated matching and *β* within a simulated admixed population. **A**: ‘deep’ simulation with population splits A–B and (AB)–C of 10^4^ and 4 *×* 10^4^ generations respectively. **B**: ‘shallow’ simulation with population splits A–B and (AB)–C of 10^3^ and 4 *×* 10^3^ generations respectively. *β* coefficients were calculated from matching coefficients following equation 1. Color indicates the mean sub-population contribution to each pair of genotypes with red, green and blue channels respectively corresponding to sub-populations A, B and C.

Visualization of pair-wise matching estimates from the deep simulation revealed a pair of triangular patterns relating to estimates between pairs of distinct individuals and estimates of self-matching (i.e., diagonal values of the matching coefficient matrix). The approximate bounds of estimates between pairs of distinct individuals corresponded to matching between (1) individuals within the same sub-population; (2) between individuals of sub-populations A–B; and (3) between individuals of sub-populations (AB)–C. Estimates involving admixed individuals were scattered throughout the triangle in rough accordance with mean sub-population membership of each pair of individuals. The few outliers observed to the lower right of this triangular pattern were associated with relationships between pairs of individuals that contained an unusually closely related haplotype (not shown). A similar pattern was observed for self-matching coefficient estimates in which a rough triangular shape was bounded by the self-matching coefficients of (1) non-admixed individuals; (2) admixed-individuals descending from sub-population A–B; and (3) admixed individuals descending from (AB)–C. Self-matching and pair-matching estimates were clearly separable when calculated from microhaplotype markers but showed substantial overlap when estimated from the equivalent SNV markers.

The double triangle pattern of the deep simulation was not immediately evident within the shallow simulation although substantial rank order variation was present as evidenced by the apparent jitter between estimation methods (Figure 6B). Larger matching coefficients were observed for pairs of individuals within a single sub-population (pure red, green, or blue points within Figure 6B) and the smallest matching coefficients were observed between pairs of individuals from sub-population (AB)–C. Self-matching estimates were clearly distinct from both SNV- and microhaplotype-based methods.

The mean squared error of microhaplotype estimates was less than that of SNV-based estimates across all tested reference population points (Table 3). The only exception to this trend occurred with a reference point of 10^4^ generations that greatly exceeded the deepest sub-population split of 4*×*10^3^ generations (Table 3B). The MSEs of both SNV and microhaplotype estimates were observed to decrease with respect to reference population depth, with the exception of the deepest reference point in the case of microhaplotype estimates.

**Table 3.** Mean squared error of simulation estimates with respect to proportion of shared IBD segments. **A**: ‘deep’ simulation with population splits A–B and (AB)–C of 10^4^ and 4 *×* 10^4^ generations respectively. **B**: ‘shallow’ simulation with population splits A–B and (AB)–C of 10^3^ and 4 *×* 10^3^ generations, respectively.

| A |  |  | B |  |  |
| --- | --- | --- | --- | --- | --- |
| $\delta_{\text{ref}}$ | $\text{MSE}[\hat{\beta}^s]$ | $\text{MSE}[\hat{\beta}^h]$ | $\delta_{\text{ref}}$ | $\text{MSE}[\hat{\beta}^s]$ | $\text{MSE}[\hat{\beta}^h]$ |
| $10^0$ | 0.1075 | 0.0071 | $10^0$ | 0.0078 | 0.0033 |
| $10^1$ | 0.1075 | 0.0071 | $10^1$ | 0.0077 | 0.0033 |
| $10^2$ | 0.1065 | 0.0067 | $10^2$ | 0.0075 | 0.0031 |
| $10^3$ | 0.0973 | 0.0039 | $10^3$ | 0.0055 | 0.0019 |
| $10^4$ | 0.0534 | 0.0105 | $10^4$ | 0.0004 | 0.0021 |

### Kiwifruit germplasm data

Identification of Capture-bait sites yielded a BED file of 7018 haplotype assembly targets. The full haplotype calling pipeline was split into 10 processes by splitting the BED file into chunks of 702 loci (700 in the final chunk). Each chunk ran with an average wall-clock time of approximately 15.25 hours (standard deviation of 2.25 hours) and 2.16 GB of RAM (standard deviation of 0.17 GB). We note that computational resource utilization is reported from the job scheduler of a production cluster with other workloads running concurrently, and values should be considered as approximate indications only. A total of 6218 polymorphic haplotype loci were retained after filtering with sgkit. The number of haplotype alleles per retained assembly locus ranged from 2 to 103, with a mean of 35.6 (Figure 7A). The number of SNVs within each haplotype locus was approximately normally distributed with a mean of 29.0 and standard deviation of 9.2 (Figure 7B).

**Fig. 7.**
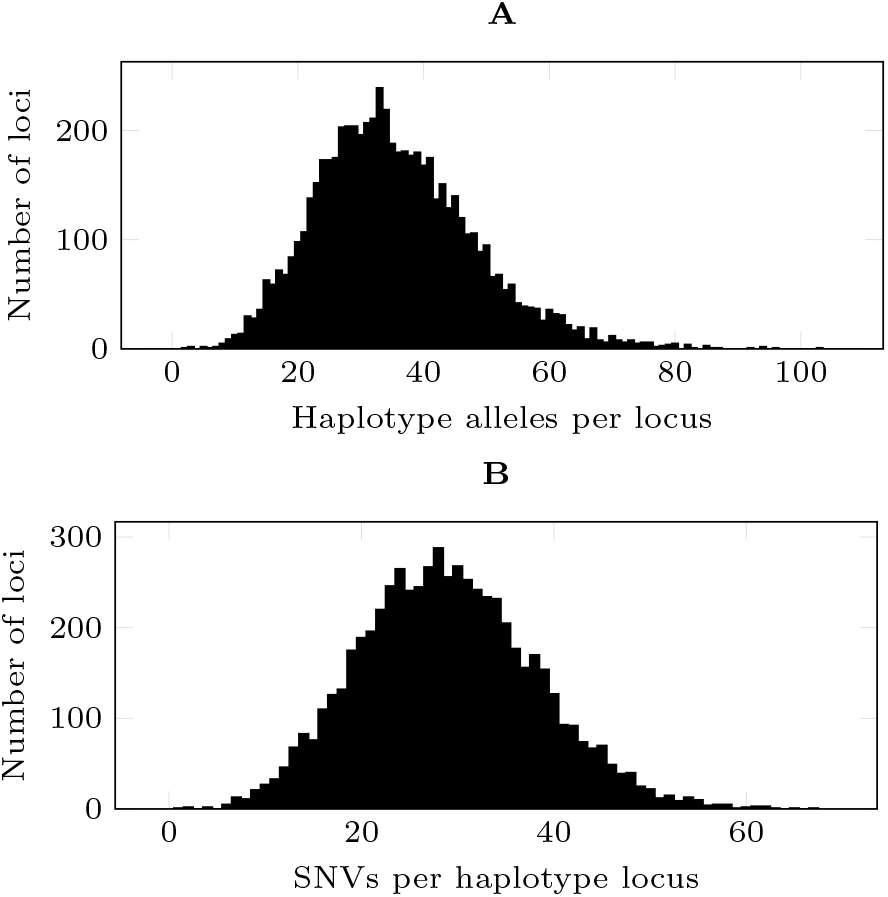
Allelic variation within 100bp assembly loci. **A**: Observed haplotype alleles per assembly locus. **B**: Observed single nucleotide variants (SNVs) per assembly locus.

Principal component analysis of microhaplotype and SNV matching coefficients produced broadly similar patterns of population structure (Figure 8), with a substantially higher proportion of variance being explained by the first component of the SNV analysis (79.7%) than that of the microhaplotype analysis (60.4%). Members of the AC, CC and PC clades were clearly distinguishable based upon the first two components in both analyses, although greater distinction was observed in the analysis of SNV data (Figure 8C). Within the PC clade, the two *A. valvata* samples and single *A. macrosperma* sample clustered closely with the OH clade in the microhaplotype analysis, and to a lesser degree in the SNV analysis (*v* and *m* in Figure 8). Members of the RC clade generally fell between the CC and OH clades, with the single *A. chinensis* var. *setosa* sample grouping closely to *A. chinensis* var. *deliciosa* (CC) in the microhaplotype analysis and within the *A. chinensis* var. *deliciosa* cluster in the SNV analysis (*s* and *d* in Figure 8). Component three of both analyses separated the OH clade with *A. rufa* (RC), clustering closely in the microhaplotype analysis (*r* in Figure 8B) and to a lesser degree in the SNV analysis (Figure 8D). Individuals of the CC clade generally clustered according to ploidy, including *A. chinensis* var. *chlorocarpa* and *A. chinensis* var. *coloris* samples grouping with *A. chinensis var. deliciosa* samples. Within the AC clade, tetraploid samples of *A. arguta* var. *arguta* and *A. arguta* var. *purpurea* tended to group together with higher ploidy samples separated. However, diploid and tetraploid samples of *A. melanandra* grouped together.

**Fig. 8.**
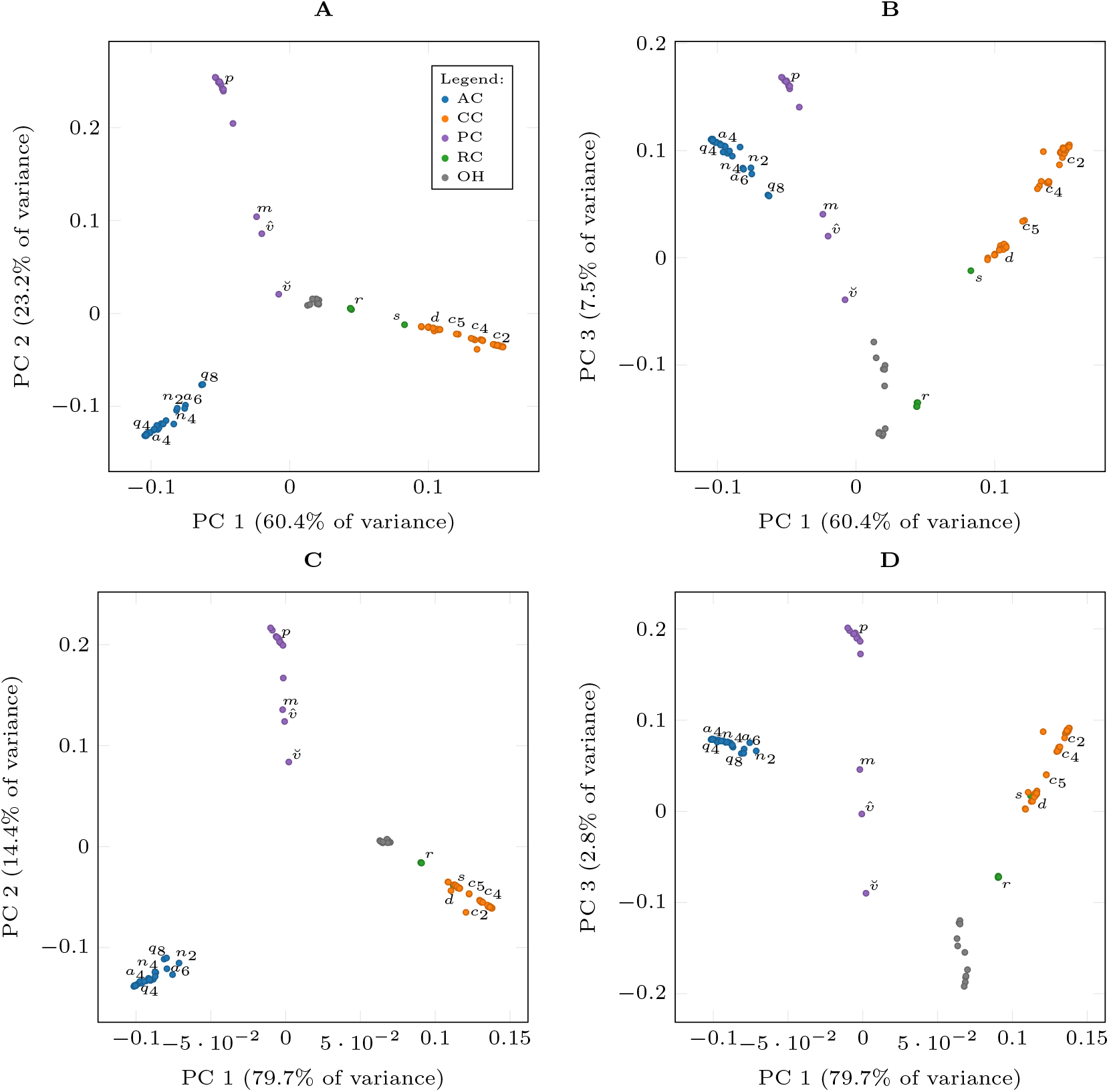
Principal components analysis of matching coefficients **A** and **B**: estimated from microhaplotypes. **C** and **D**: estimated from single nucleotide variants (SNVs). Colors indicate taxonomic clades following Table 2 as defined in the legend of **A**. Letters indicate the approximate mean position of sample within specific taxa with *a*: *Actinidia arguta* var. *arguta, c*: *A. chinensis* var. *chinensis, d*: *A. chinensis* var. *deliciosa, s*: *A. chinensis* var. *setosa, r*: *A. rufa, m*: *A. macrosperma, n*: *A. melanandra, p*: *A. polygama*, and *q*: *A. arguta* var. *purpurea*. Male and female samples of *A. valvata* are respectively indicated with 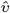 and 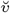. Ploidy is indicated by subscript were it varies within taxon. Note that samples of *A. chinensis* var. *chlorocarpa* and *A. chinensis* var. *coloris* group within the *A. chinensis* var. *deliciosa* samples. Symbols have been adjusted to minimize over-plotting.

Visual comparison of the matching and *β* coefficients estimated from microhaplotype- and SNV-encoded variant data yielded a curved pattern consistent with theoretical expectations (Figure 9). However, this curved pattern was not immediately obvious from relationships among individuals within any single one of broadly defined taxonomic clades (Table 2). Rank-order variation in relationships between SNV- and microhaplotype-based estimates were abundant within each of the broadly defined clades. There was clear separation between relationships of the three best represented clades (AC, CC and PC). Notably, genotype pairs within CC exhibited greater SNV matching than those of AC given a similar degree of microhaplotype matching. The individual with the highest self-matching coefficient was the single sample of *A. chinensis* var. *setosa*. This sample also exhibited higher matching coefficients with other members of *A. chinensis* in the CC clade than with *A. rufa* of its nominal clade (RC).

**Fig. 9.**
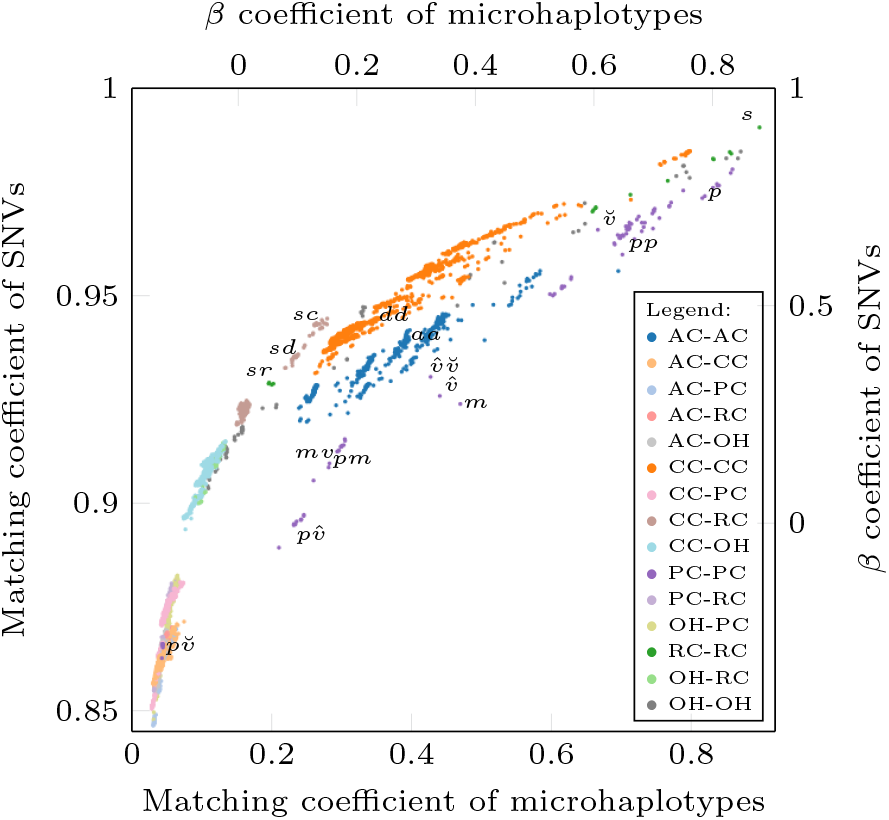
Estimated matching and *β* coefficients between *Actinidia* germplasm samples. Matching coefficients were estimated from microhaplotype calls encoded as microhaplotypes and SNVs following equation 2, as implemented in sgkit. *β* coefficients were calculated from matching coefficients following equation 1. Colors indicate relationships within or between taxonomic clades following Table 2 as defined in the legend. Symbols indicate approximate (mean) positions of selected taxon self-matching (single letter) and pair-matching (pairs of letters) with *a*: *A. arguta* var. *arguta* (tetraploid), *c*: *A. chinensis* var. *chinensis, d*: *A. chinensis* var. *deliciosa, m*: *A. macrosperma, p*: *A. polygama, r*: *A. rufa, s*: *A. chinensis* var. *setosa*, and *v*: *A. valvata* (with 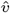 and 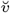 indicating the male of female sample respectively). Symbols have been adjusted to minimize over-plotting, with the exceptions of *aa* and *dd*.

Matching coefficients among *A. polygama* samples were generally higher than those of other taxa when estimated from microhaplotype markers, but comparable to single taxon matching coefficients of taxa within CC when estimated from SNV markers (Figure 9). Matching coefficients between *A. polygama* and *A. macrosperma* were relatively low (0.913) when estimated from SNVs but comparable to other within-clade coefficients (0.295) when estimated from microhaplotypes. The substantial dissimilarity between male and female samples of *A. valvata* observed in Figures 8 and 9 reflected differences in their matching coefficients with other members of the PC clade as shown in Table 4. Both the *A. macrosperma* and male *A. valvata* samples had relatively low self-matching coefficients when estimated from microhaplotypes (0.469 and 0.440, respectively), or SNVs (0.924 and 0.926, respectively). The SNV-based self-matching of the male sample (0.926) was lower than the matching coefficient between the male and female *A. valvata* samples (0.930). This contrasted a lower matching coefficients (mean of 0.866) seen between the *A. polygama* and female *A. valvata* samples, which were comparable to values seen between broader taxonomic clades (Figure 9). The female *A. valvata* sample had a much higher self-matching estimated from microhaplotypes or SNVs (0.666 and 0.966, respectively) than its male counterpart, although lower than that observed for many other taxa, including *A. polygama* (0.816 and 0.974, respectively).

**Table 4.** Matching coefficients within PC. **A**: estimated from microhaplotypes. **B**: estimated from single nucleotide variants (SNVs). 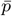. *polygama* samples. *m*: *A. macrosperma* sample. 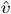. *valvata* sample. 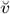. *valvata* sample.

| A |  |  |  |  |
| --- | --- | --- | --- | --- |
| | $\bar{p}$ | *m* | $\hat{v}$ | $\check{v}$ |
| $\bar{p}$ | 0.816 | 0.295 | 0.236 | 0.044 |
| *m* | 0.295 | 0.469 | 0.282 | 0.281 |
| $\hat{v}$ | 0.236 | 0.282 | 0.440 | 0.427 |
| $\check{v}$ | 0.044 | 0.281 | 0.427 | 0.666 |

| B |  |  |  |  |
| --- | --- | --- | --- | --- |
| | $\bar{p}$ | *m* | $\hat{v}$ | $\check{v}$ |
| $\bar{p}$ | 0.974 | 0.913 | 0.895 | 0.866 |
| *m* | 0.913 | 0.924 | 0.910 | 0.909 |
| $\hat{v}$ | 0.895 | 0.910 | 0.926 | 0.930 |
| $\check{v}$ | 0.866 | 0.909 | 0.930 | 0.966 |

Estimates of haplotype and nucleotide diversity within well-represented taxon–ploidy combinations (Table 5) were consistent with observations made from Figure 9. *A. polygama* exhibited the lowest diversity among the included taxa. *A. chinensis* var. *deliciosa* had the highest haplotype diversity and *A. arguta* var. *arguta* the highest nucleotide diversity. Notable, the inversion in rank-order of gene and nucleotide diversity between *A. chinensis* var. *deliciosa* and *A. arguta* var. *arguta* was consistent with their observed positions in Figure 9.

**Table 5.**
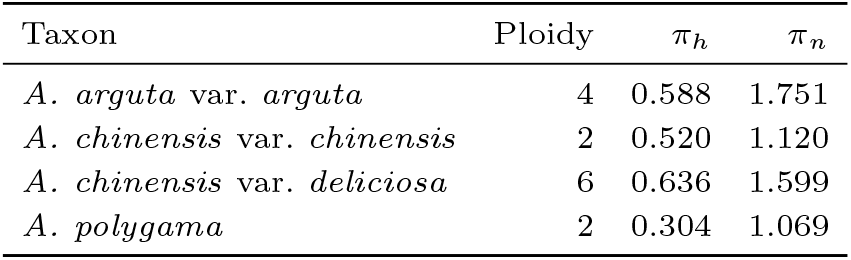
Observed molecular diversity of *Actinidia* taxa with ten or more samples. *π*_*h*_ is the genome-wide mean of haplotype diversity (Nei, 1973) and *π*_*n*_ is the genome-wide mean of nucleotide diversity (Nei and Li, 1979) within haplotype loci.

## Discussion

Despite extensive interest in applying haplotype markers to complex autopolyploid crops, there has been little examination into the impact of microhaplotype markers on kinship estimation in similarly complex taxa. Recently, Vexler et al. (2025) proposed that haplotype-based estimates of relatedness will emphasize recent demographic history. Our findings corroborate this by building upon the work of Weir and Goudet (2017) and Bilton (2020), and developing theoretical expectations for kinship estimates from microhaplotype markers. Our findings have clear parallels to the existing literature on the importance of marker size including runs of homozygosity (McQuillan et al., 2008) and the distinction between gene and nucleotide diversities (Nei, 1973; Nei and Li, 1979). Whilst our theoretical expectations were developed in terms of the matching and *β* coefficients, they are equally applicable to any kinship estimates found by scaling matching or *β* to a relevant reference population. A further potential benefit of microhaplotype markers is improved dosage accuracy in autopolyploids, which would also improve kinship estimation. Dosage accuracy is not explored within our analyses, as our SNV calls were found via atomizing haplotypes, thereby guaranteeing equivalent dosages of microhaplotype and SNV markers.

Within this research, we also developed Bayesian methods for sequence-based microhaplotype assembly in autopolyploids and we make these methods available through the MCHap software package. The MCHap package includes methods for both the *de novo* assembly of microhaplotypes and genotype calling from a set known microhaplotypes. Our methods of microhaplotype assembly are suitable for extremely diverse populations of variable ploidy as evidenced by their application in a germplasm selection spanning 17 taxa and 5 ploidy levels.

### Coalescent simulations

Our coalescent simulations assume an extremely simple demographic history with somewhat arbitrary evolutionary parameters. These simulations are not intended as hypotheses of evolutionary history, but rather as a means to confirm theoretical expectations of kinship estimation from SNVs and microhaplotypes. The congruence between theoretical expectations and simulation results is most apparent when observing that the bounding points of either triangular pattern in Figure 6A are analogues of the points *μ*_1_, *μ*_3_ and *μ*_7_ in Figure 3. These simulations corroborate our theoretical expectations that (a) kinship estimation from microhaplotypes reduces the influence of evolutionary distant alleles; (b) this results in a more accurate estimate of kinship with respect to a recent reference point (or population); and (c) these differences can result in dramatic changes in the rank-order of relationships with respect to SNV-based estimates.

The observed increase in mean squared error of microhaplotype estimates with respect to the deepest reference point (Table 3) is contrary to the expected monotonic decrease in variance with increased depth of the reference population (equation 11). The increased MSE is instead explained by limitations in calculation of ‘true’ kinship. This calculation considered only inheritance of chromosomal segments and disregarded mutations with respect to the ancestral allele which is not strictly in accordance with the principal of IBS as a precondition of IBD (Cotterman, 1940; Malécot, 1969; Jacquard, 1972). Incorporating mutations into this calculation would require the specification of a minimum allele length for determining the non-mutation of ‘alleles’, as is done in the analysis of runs of homozygosity (McQuillan et al., 2008). The choice of minimum allele length is subjective and a larger minimum length would probably favor microhaplotype-based estimation of kinship hence our decision to ignore mutations and consider only inheritance. We expect that the exclusion of mutations from the calculation of ‘real’ kinship would primarily disadvantage microhaplotype-based estimates of kinship which are more sensitive to changes in allelic state. Therefore, we consider the MSE estimates reported in Table 3 to be inflated, especially those of microhaplotype-based estimates with respect to deeper reference points.

### Analysis of germplasm

The *Actinida* germplasm population analyzed within this manuscript exhibits extreme allelic diversity, with approximately 30% of base positions varying to some degree within the observed genotype calls (Figure 7B). The decision to restrict haplotype assembly to 100bp loci was in part motivated by prior knowledge of the extreme heterozygosity, and also to simplify the interpretation of results in the context of theoretical expectations. The requirement for 98% of samples having a read depth of 50 or higher was also intended to reduce the prevalence of bait targets with poor yield. Despite our conservative approach, we expect a significant rate of genotype calling error within our analyses owing to structural variation and evolutionary divergence among taxa (Cheng et al., 2025). Furthermore, the capture-sequencing array used within this analysis was originally designed from diploid *A. chinensis* var. *chinensis* data (Tahir et al., 2020) and sequences were aligned to the diploid *A. chinensis* var. *chinensis* ‘Donghong’ reference genome (Han et al., 2023). Hence, we would expect the highest rate of genotyping error to occur among members of the AC and PC clades, which are the most evolutionarily distant from *A. chinensis*. The paleo-polyploid origins of *Actinidia* (He et al., 2003, 2005) and presence of putative allopolyploid species such as *A. valvata* (Hu et al., 2025) may further reduce the accuracy of read mapping and genotype calls (Cheng et al., 2025). In spite of these concerns, we consider the broad observations borne from these data to be indicative of the real allelic variation and relationships among samples. We do, however, note that the samples used within this analysis are a subset of the New Zealand germplasm collections, and that the diversity represented by these collections may be biased with respect to natural populations. In particular, we would caution readers against assuming that the within-taxon diversity reported here is representative of natural populations.

Our analyses revealed a population structure that is in broad agreement with the treatment of Liu et al. (2017). The most distinct clusters of samples within our PCAs corresponded to the AC, CC, PC and OH clades. These groupings were more distinct in the PCA performed on matching coefficients estimated from SNV data (Figure 8C and D), which fits with the theoretical expectation that SNV-based analysis increases the weighting of deeper coalescent branches. This is further supported by the larger proportion of variance attributed to PC1 (in the SNV analysis), which corresponds to the deepest phylogenetic split (Liu et al., 2017). Interestingly, taxa and samples with lower ploidy tended to be located nearer to the extremities of the PCA which appears to correspond with slightly elevated matching coefficients among distant taxa with higher ploidy. It is possible that this occurs because of increased retention of alleles in higher ploidy populations, resulting in slightly increased matching coefficients with distant relatives. However, a similar result may arise as an artifact of increased genotype calling error in higher ploidy samples (Cooke et al., 2021). We note that the tight clustering of all three hexaploid *A. chinensis* varieties (not shown in figures because of over-plotting) is unsurprising given their past treatment as varieties of a separate species *A. deliciosa*. The placement of the two pentaploid *A. chinensis* var. *chinensis* samples between tetraploid *A. chinensis* var. *chinensis* and hexaploid *A. chinensis* var. *deliciosa* could also suggest a hybrid origin.

Visual comparison of matching coefficients estimated from SNV and microhaplotype markers highlighted differences in the dispersion of SNVs among haplotypes of the sampled AC, CC and PC clades, as evidenced by the observed separation of intra-clade relationships (Figure 9). We consider this to be indicative of distinct demographic histories among the sampled individuals of these clades resulting in distinct haplotype and nucleotide diversities (Nei, 1973; Nei and Li, 1979; Tajima, 1983). Samples from the CC clade had higher SNV-derived matching coefficients than their microhaplotype-derived estimates. This is consistent with a hypothesis of a relatively recent radiation from a small ancestral population, such that unique in-descent haplotypes have accumulated relatively few SNV mutations (Tajima, 1983). In contrast, members of the AC clade had relatively low SNV-derived matching coefficients for a similar degree of microhaplotype matching, which is suggestive of an older ancestral population. Whilst these trends are somewhat apparent from simple diversity metrics (Table 5), the broader detail of inter- and intra-clade relationships are captured by visualizing SNV and microhaplotype estimates, as in Figure 9. Samples of *A. polygama* exhibited higher microhaplotype-derived matching coefficients than either AC or CC but with proportionally low SNV-based estimates. This could support a hypothesis of an older population that has contracted in size, resulting in the loss of haplotype diversity. However, this interpretation may be biased by non-uniform sampling of the natural population.

Perhaps the most notable feature of Figure 9 is the outlying relationships between taxa of the PC clade. The most extreme of these is the relationship between the female *A. valvata* sample and samples of *A. polygama*, which is comparable to that of the most distantly related clades. A simple conclusion is that this may be an incorrectly labeled sample of some other clade. However, we did not observe any relationship between this sample and those of other clades that would support such a hypothesis. Furthermore, this sample does bear some relationship to the male *A. valvata* and *A. macrosperma* samples, which are in turn related to the *A. polygama* samples. The position of relationships involving these two samples and their nearest relatives are consistent with what we would expect for hybrid or admixed samples (reminiscent of *μ*_4*−*5_ in Figure 3). This hypothesis is also supported by the low self-matching coefficients observed for these samples (Table 4), which indicate high heterozygosity. A recently published genome assembly of *A. valvata* found compelling evidence in favor of an allopolyploid origin of the taxon (Hu et al., 2025). They suggest that an allopolyploid *A. valvata* was derived from a pair of taxa closely aligning to *A. polygama* and *A. macrosperma*. This hypothesis, in combination with some mislabeling of samples, could potentially explain our observation. However, the expansive genetic distances observed among our samples of this clade are still surprising and warrant further investigation.

Relationships observed between clades indicated that CC, RC, and OH are more closely related to one another than to AC or PC, in agreement with prior phylogenic studies. However, our results suggest a closer relationship between CC and RC than between RC and OH (Figure 9) which conflicts with the phylogeny of Liu et al. (2017). Furthermore, we found a closer relationship between *A. chinensis* var. *setosa* and other *A. chinensis* varieties than with with *A. rufa*. The matching coefficients between *A. chinensis* var. *setosa* and other *A. chinensis* varieties were indicative of low SNV differentiation among haplotypes, which would be consistent with a hypothesis of recent isolation. But these observations should be interpreted with care, given that are drawn from a single sample of *A. chinensis* var. *setosa*.

### Relevance to crop breeding

The development of tools suitable for the genetic analysis of polyploid plant species has lagged behind those of their diploid counterparts, despite the global importance of polyploid crops (Bourke et al., 2018; Cheng et al., 2025). A key difficulty in the analysis of polyploid plant species is the application of variant calling tools, which have typically been developed for less heterozygous, diploid species. A common approach in variant calling software is the direct enumeration of all possible genotypes given a set of putative alleles (see equation 13). This approach scales poorly with both ploidy and heterozygosity, and discourages the analysis of more complex variants. We argue that probabilistic methods such as the Gibbs-sampler are a better fit in autopolyploid crops because they make calling complex variants tractable and, in doing so, can turn heterozygosity into an advantage. Furthermore, the use of complex microhaplotype markers is a natural fit with targeted sequencing technologies, which are rapidly being adopted in autopolyploid breeding programs (Tahir et al., 2020; Zhao et al., 2023; Clare et al., 2024; Endelman et al., 2024; Zhao et al., 2024b,Within this manuscript we have chosen to demonstrate the utility of microhaplotypes within a diverse *Actinidia* germplasm collection. The motivation for this choice was two-fold: (1) to emphasize that our methods are tractable in extremely heterozygous populations; and (2) to confirm our theoretical expectations over a wide range of evolutionary distances. However, we also consider our methods to be applicable within advanced selections as long as sequencing targets regularly contain multiple basis SNVs for assembly, as is the case in *Actinidia*. We note that the branch lengths of the coalescent are expected to be highly variable even under simple demographic histories (Degnan and Salter, 2005), which will result in the uneven distribution of SNVs among haplotypes. We would expect our methods to be particularly relevant in breeding programs which utilize wide hybrids to introgress new genetic variation owing to the higher SNV weighting of the introgressed alleles. If uncounted for, an uneven distribution of SNVs among the alleles of a population may degrade the signal of recent population structure. Powell et al. (2010) argue that the principal use of the IBD concept within quantitative genetics is to infer IBS at unobserved loci via linkage. In light of this, we would expect IBD estimated with respect to a recent reference population to be more informative than those of a deeper reference point. Here, we have demonstrated that microhaplotype markers are a practical means of achieving this goal in complex heterozygous taxa.

## Data availability

The raw sequencing data supporting the findings of this article are available in the NCBI Sequence Read Archive under BioProject accession PRJNA1505659.

## Author contributions statement

T.M. experimental design, software, theory and writing. E.K. selection, processing of germplasm samples and writing. A.H. bioinformatics processing and writing. A.G. selection and processing of germplasm samples. S.T., J.M. P.W. and M.B. supervision and manuscript review.

## Acknowledgments

The authors acknowledge Paul Datson and Timothy Bilton for their valuable feedback on an earlier draft of this manuscript.

