## Supplemental Text 1 for "Microhaplotypes Improve Kinship Estimation in Heterozygous, Mixed-Ploidy Populations of *Actinidia*"

### S1 The effect of monomorphic loci on the matching and $\beta$ coefficients

Within our research we first define an estimate of the matching coefficient relative to the number of polymorphic loci in the sample-population as

$$\hat{m}^s_{GG'} = \frac{1}{kk'} \sum_{i \in G} \sum_{j \in G'} \frac{s_{\text{pop}} - s_{ij}}{s_{\text{pop}}} \tag{1}$$

where $s_{\text{pop}}$ is the total number of SNVs (polymorphic sites) found in the sample population, $s_{ij}$ is the number of SNVs differentiating $i^{\text{th}}$ haplotype of $G$ from $j^{\text{th}}$ haplotype of $G'$, and $k$ and $k'$ are the ploidy of $G$ and $G'$ respectively.

Equation 1 is equivalent to the formulation given in Table 2 of Weir and Goudet (2017) which assumes diploidy

$$\tilde{M}^i_{jj'} = \frac{1}{4} \sum_u X^i_{ju} X^i_{j'u} \tag{2}$$

where $X^i_{ju}$ is the dosage of the $u^{\text{th}}$ allele in the $j^{\text{th}}$ inidividual of sub-population $i$ (sub-population is ignored in our work). An important distinction between equations 1 and 2 is that in equation 1 we sum over allele copies (combinations of alleles drawn from $G$ and $G'$) where as in equation 2 Weir and Goudet sum over allelic states $(0, ..., u)$. The formulation of Weir and Goudet (2017) is given for a single locus and hence does not take a mean over multiple loci. Our formulation in equation 1 accounts for multiple loci by the use of the $s_{pop}$ and $s_{ij}$ terms. This is an intentional choice to both link the definition to the variations between individual haplotypes ($s_{ij}$) and to SNV discovery within a sample-population ($s_{\text{pop}}$).

We later redefine the matching coefficient in terms of some genomic length/interval of length $n$ from which alleles have been discovered as

$$\hat{m}^s_{GG'} = \frac{1}{kk'} \sum_{i \in G} \sum_{j \in G'} \frac{n - s_{ij}}{n} \quad \text{where } n \geq s_{ij}. \tag{3}$$

Equation 3 results in a larger matching coefficient because $n$ includes monomorphic loci in addition to the polymorphic loci identified as $s_{\text{pop}}$. However, this does not affect the estimation of $\beta$ because the value of $s_{\text{pop}}$ or $n$ is factored out of the calculation as follows

$$\begin{aligned}
\hat{m}^s_{GG'} &= \frac{1}{kk'} \sum_{i \in G} \sum_{j \in G'} \frac{n - s_{ij}}{n} \\
&= \frac{n - \frac{1}{kk'} \sum_{i \in G} \sum_{j \in G'} s_{ij}}{n} \\
&= \frac{n - p}{n} \quad \text{where } p = \frac{1}{kk'} \sum_{i \in G} \sum_{j \in G'} s_{ij} \\
&= 1 - \frac{p}{n}.
\end{aligned}$$

The equivalent for the population mean is

$$\tilde{m} = 1 - \frac{\tilde{p}}{n}.$$

Now substitute these terms into the estimator of $\beta$

$$\begin{aligned}
\hat{\beta}_{GG'} &= \frac{m_{GG'} - \tilde{m}}{1 - \tilde{m}} \\
&= \frac{\left(1 - \frac{p}{n}\right) - \left(1 - \frac{\tilde{p}}{n}\right)}{1 - \left(1 - \frac{\tilde{p}}{n}\right)} \\
&= \frac{\frac{\tilde{p} - p}{n}}{\frac{\tilde{p}}{n}} \\
&= \frac{\tilde{p} - p}{\tilde{p}}
\end{aligned}$$

which does not depend upon $n$ or, in the alternative formulation, $s_{\text{pop}}$.
